# Active eye modeling: a tool for estimating retinal image quality across the visual field in everyday life

**DOI:** 10.64898/2026.11.04.764163

**Authors:** Charlie S. Burlingham, Ian M. Erkelens, Daniel P. Spiegel, Aaron Nicholls, Oliver S. Cossairt, Phillip Guan

**Author notes:** **Funding:** This work was funded by Meta Platforms, Inc. No external funding was received. **Commercial Relationships:** C.S. Burlingham, Meta (E); I.M. Erkelens, Meta (E); D.P. Spiegel, Meta (E); A. Nicholls, Meta (E); O.S. Cossairt, Meta (E); P. Guan, Meta (E).

## Abstract

**Purpose:** The “visual diet,” the totality of visual experience, shapes retinal image formation and thereby refractive development, but measuring retinal image statistics during everyday behavior is challenging. Limits in mobile eye tracking, egocentric video, and full-field optical measurement leave gaps in our understanding of how environmental and behavioral factors shape image quality across the retina.

**Methods:** We introduce active eye modeling, a computational platform combining optically validated, wide-field, accommodating eye models^1^ with realistic three-dimensional virtual environments and gaze behavior derived from human data. This approach enables systematic estimation of retinal blur across the visual field under parametrically controlled conditions, including varying viewing distances, ambient light levels, and environmental depth structure. Inputs may be simulated and/or drawn from real-world measurements: scene depth, video, and gaze/pupil recordings.

**Results:** During simulated far work, peripheral image quality is depth-tolerant, owing to wide depth-of-field from off-axis aberrations. Near work under dim indoor lighting substantially increases peripheral blur, reducing contrast at mid and high spatial frequencies. This contrast reduction is comparable in magnitude to a moderate diffusing filter (Bangerter 0.4 strength) that affects eye growth when imposed experimentally in animal models. A consistency check with eye-tracked virtual driving confirms peripheral depth-tolerance during far work.

**Conclusions:** Active eye modeling provides a flexible framework for investigating how visual experience shapes the retinal image, relevant to refractive development, display and lens design, and peripheral vision. Applied to common visual behaviors, the platform identifies optical conditions during indoor near work that warrant empirical investigation, complementing measurement-based approaches.

## 1 Introduction

Myopia prevalence is increasing globally, with substantial geographic and generational variation that points to environmental rather than genetic drivers.^2,3^ Rates have reached 91% in Singapore and 42% in the United States, where prevalence has roughly doubled over three decades, yet remain 17% in Mongolia and 6% in parts of Africa.^2,4^ Among the environmental factors implicated in this increase, two stand out as the most robustly supported by current evidence: insufficient time spent outdoors^5,6^ and excessive near work.^5,7,8^ The “visual diet,” the totality of visual experience during development, encompassing viewing distances, light levels, environmental structure, and visuo-motor behavior, has changed substantially with industrialization across much of the world.^2,9–11^ Understanding how these changes affect the developing eye requires tools that can systematically characterize the optical consequences of everyday behavior.

The developing eye starts out, on average, hyperopic and elongates during emmetropization until the retinal image is well-focused, at which point growth rate decreases.^12^ This process operates as a closed-loop feedback system, i.e., that “the function of emmetropization is to minimize blur^3^,” or, relatedly, to maximize retinal image contrast. The tight distribution of refractive errors in non-myopic populations attests to the precision of this process. Multiple interacting signals drive emmetropization, including directional cues that encode the sign of defocus and non-directional cues related to overall image quality.^3,12^

Animal studies have documented that a wide range of optical interventions can alter eye growth: form deprivation, lens-induced defocus, narrow-band spectral illumination, and peripheral-only stimulation each reliably modulate axial length and refraction,^13–17^ with effects that can be local to specific retinal regions and that persist into adolescence^3,15,16,18,19^. How the eye determines the sign of defocus from the retinal image remains an area of active investigation, with proposed mechanisms including computations based on higher-order aberrations, longitudinal chromatic aberration, and vergence-accommodation coupling.^3,20–24^ Progress on these directional cues has renewed interest in how the eye’s optics interact with the visual diet during development.^2,11,25^

One directional hypothesis posits that near work induces relative peripheral hyperopic defocus, causing the developing eye to elongate to bring the peripheral image into focus.^26^ However, the relationship between peripheral defocus and eye growth is more complex than this intuition suggests: the actual peripheral defocus pattern depends on the fixation point, accommodative state, and the full dioptric landscape of the environment. Ophthalmic interventions that modify peripheral optics, such as orthokeratology, were motivated primarily by animal studies demonstrating that imposed peripheral defocus can modulate eye growth^15,16^, and have proven effective in slowing myopia progression.^10^ Yet the finding that lenses inducing the opposite sign of peripheral defocus also slow progression^10,27^ raises important questions about whether defocus direction alone is the critical variable.

Non-directional cues also play an established role in eye growth regulation.^24^ In form deprivation, diffusing filters degrade retinal image contrast and phase structure, reliably producing myopia in animal models^24,28^, even when restricted to the peripheral retina.^15^ Contrast signaling through retinal ON and OFF pathways may also contribute to refractive development, with disruptions in these pathways associated with myopic shifts.^29,30^ Mild peripheral contrast reduction through Diffusion Optics Technology (DOT) lenses slows myopia progression,^29,31^ whereas the stronger contrast reduction imposed in form deprivation induces myopia. This apparent contradiction suggests that contrast magnitude alone does not determine the growth response, and that factors such as spatial frequency content, temporal dynamics, or accommodative interactions also contribute. These apparent contradictions underscore the need for tools that can estimate actual retinal image statistics during everyday viewing.

There is strong evidence that time spent outdoors is protective against myopia,^6,32^ though the mediating mechanisms remain debated. Candidate factors include increased retinal illumination, further viewing distances, spectral characteristics of daylight, higher spatial frequency content in natural scenes,^33,34^ and increased physical activity.^6,11^ A leading mechanistic explanation is that bright light stimulates retinal dopamine release, which inhibits excessive eye growth — clearly demonstrated in animal models.^9,35–38^ The likely interaction of multiple factors motivates computational tools that can parametrically isolate individual contributions to retinal image formation.

A recognized gap lies between measurements of ocular optics and the goal of understanding how visual experience shapes refractive development.^11,25,39^ Longitudinal studies have found that relative peripheral refraction does not clearly predict who will develop myopia^26,40,41^ and may reflect a consequence rather than a cause of refractive error.^26,39^ Clinical wavefront measurements provide only a static snapshot, typically at fixation and a few peripheral locations, for an unaccommodated eye at optical infinity, whereas real-world experience involves three-dimensional environments, varying light levels, and diverse accommodative demands that jointly determine retinal image quality across the full visual field.^8,25^ Camera field-of-view and eye-box limitations in mobile eye trackers further restrict analyses mostly to central vision,^39,42,43^ excluding much of the peripheral retina that is specifically of interest in theories of eye growth.

We introduce active eye modeling as a computational platform to address this gap. Active eye models combine recently developed, highly accurate, wide-field, accommodating eye models^1^ with realistic three-dimensional virtual environments^44^ and naturalistic gaze patterns constrained by human data.^42,45^ Unlike static optical models or the geometric approximation of defocus, active eye models adaptively adjust accommodation, pupil size, and gaze direction in response to three-dimensional environmental demands, generating full-field estimates of peripheral blur and contrast that account for all ocular aberrations simultaneously^1^ across the visual field (80° *×* 50°). The same framework operates on real-world inputs: scene depth maps from camera or time-of-flight sensors, measured gaze and vergence depth, and video that can be optically degraded, allowing reconstruction of the retinal image during everyday behavior rather than only in simulated environments. More broadly, because the retinal image is the common input to all downstream vision, the platform is a general tool for any problem that depends on egocentric visual statistics, from the design of displays, spectacles, and contact lenses to naturalistic psychophysics. To demonstrate it, we simulate common indoor visual behaviors, i.e., near work and far work at varying light levels, and characterize how these conditions differentially affect the retinal image, generating testable hypotheses about how the visual diet may influence refractive development.

## 2 Methods

### 2.1 Virtual Environment

To simulate the common daily experience in post-industrial countries of indoor near and far work, we used a detailed, to-scale architectural visualization of an apartment environment. The 3D apartment scene is a commercially available asset that we modified with the 3D modelling and animation software Blender.^46^ Textures and lighting were realistic and led to a high degree of spatial detail in each rendered object. We developed custom Blender scripts to control observer eye and head movements with respect to specific environmental objects in our active observer model.

### 2.2 Active Observer Model

Informed by the average male human height (1.7 m), we positioned the virtual observer’s (single) eye at 10 hand-selected locations in the virtual apartment that were realistic, including lying in bed, sitting in bed, kneeling near a low bookshelf, sitting on a couch, standing in front of an open closet, bending down slightly to look at a rack of clothes, and walking up the stairs (Fig. 1A-B). For each viewpoint, a gaze fixation was generated by a deterministic approximation of randomly sampling an empirical distribution of fixation depths during far work (Fig. 1C) and finding the object closest to that fixation depth in the scene out of 297 observable total objects. The ray casting and intersection method used to find the object closest to the specified fixation depth took care of occlusion errors. The discrete set we evenly selected fixation depths from was (0.25, 0.4, 0.5, 0.75, 1, 1.25, 1.5, 1.75, 2, 2.25, 2.5, 3, 3.5, 4, 4.5, 5, 6 m), so for each observer position all 17 fixation depths were referenced to find an object near that depth. Once an object was fixated, we simulated a linear translation of the head and corresponding rotational tracking eye movement (translational vestibular ocular reflex, tVOR). This sort of eye-head coordination is extremely common in natural behavior, and occurs even while one is primarily stationary due to small head movements.^45^ The head movement was a straight path of 50 cm, in a random direction sampled from a 3D uniform distribution of angles. The retinal image changed throughout the tracking eye movement, and we saved out 10 linearly spaced RGB images from the point of view of the eye and corresponding retinal depth maps (Fig. 1D) with a 80° by 50° field of view (400 *×* 250 pixels). We also saved out all eye 3D positions and fixation depths. This yielded a total of 1360 unique views of the 3D scene (Fig. 1B).

**Figure 1:**
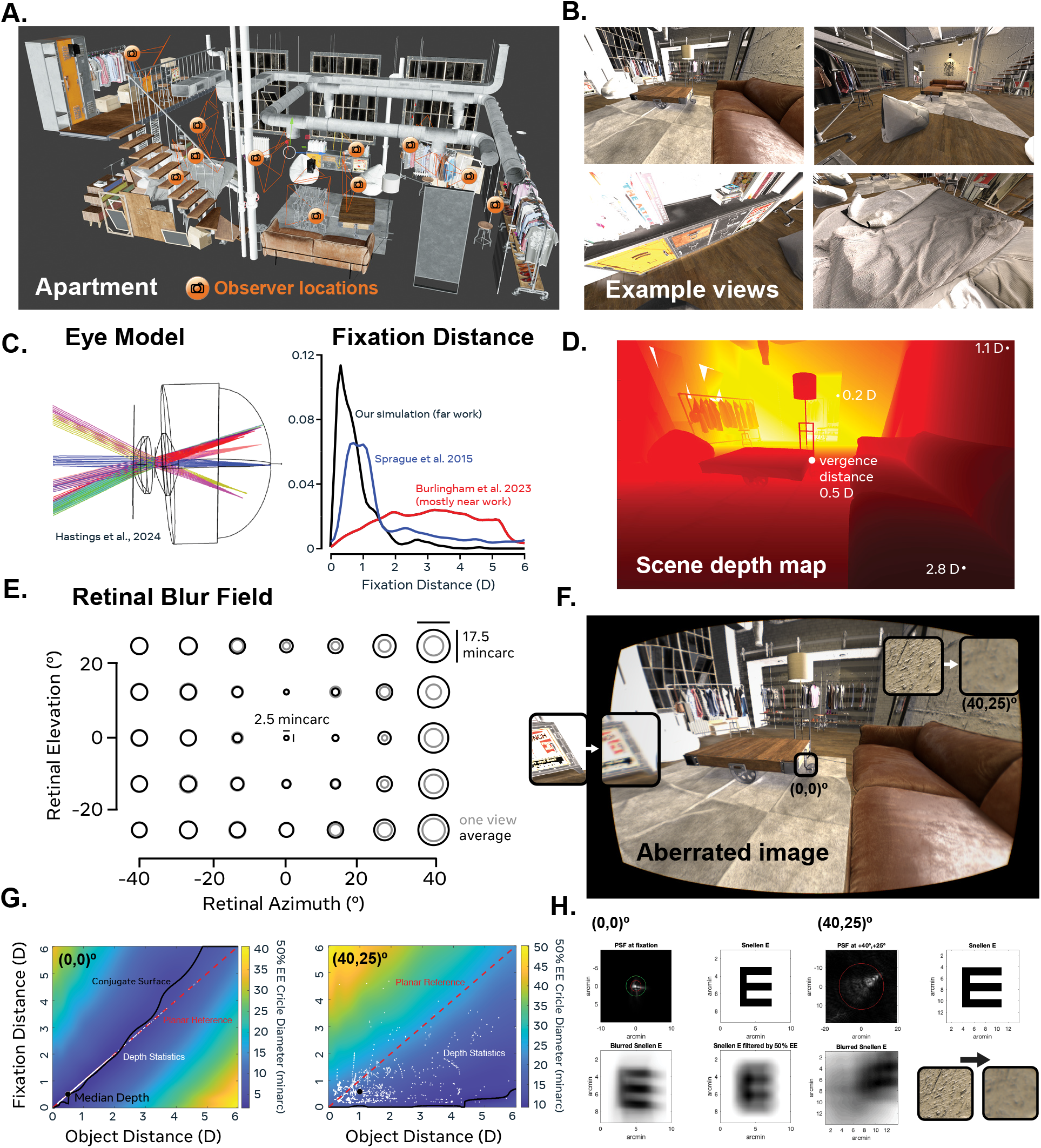
Active eye model simulation platform and estimates of the natural statistics of blur across the retina. **A.** Virtual environment, to-scale architectural visualization of an apartment scene. Orange camera icons and frustra indicate the 10 locations at which the observer was situated within the scene. Head orientation and fixation depth varied. Walls stripped for visualization purposes only. **B.** Example retinocentric views of the scene from four different observer locations. **C.** Left panel, wireframe view of typical emmetrope eye model,^1^ with light converging at various field locations. Right panel, distribution of fixation distances (convergence distances) in diopters. Black, our active observer simulation of far work in the apartment. Blue, estimates from Sprague et al.^42^ across four everyday tasks (ordering coffee, making a sandwich, walking inside, and walking outside), which included a mix of both near and far work. Red, estimates from Burlingham et.^45^ across nine everyday tasks that mostly included near work on mobile devices and computers alongside the main activity, capturing aspects of modern-day visual behavior. **D.** Depth map from one example view of the apartment, with the convergence distance at 0.5 D and other reference object depths marked in diopters. **E.** Retinal blur fields, with circles containing 50% of the energy in the PSF at each field location, blown up by a factor of 30 for visualization purposes. Grey, blur field for one view of the apartment (the example view in D). Black, average across all 3D views in the far work simulation. **F.** Aberrated image simulation, view of apartment through eye model during far work, with fixation on edge of table. Insets depict blown-up views of original RGB image and aberrated image, at (−25,0)° and (40,25)° field positions. **G.** Depth-of-field of the eye model vs. simulated depth statistics at two field positions: fixation, (0,0)° and the far periphery, (40,25)°. Colormap represents the blur circle diameter (50% encircled energy) in minarc for different combinations of fixation distance and object distance at a specific retinal location. White points, joint statistics of fixation distance and object distance projecting to that retinal location. Black circle, median fixation and object depth, for informative purposes (not used in simulation). Red dashed line, reference position of a plane perpendicular to the line-of-sight. Black solid line, retinal conjugate surface, i.e., object depth that minimizes the blur circle for each fixation distance. For (0,0)° the conjugate surface does not lie on the planar reference because of lags and leads of accommodation. **H.** Point-spread function (PSF) for 555 nm light and aberrated optotypes at two retinal locations. Upper left, PSF overlaid with circles capturing 50% (red) and 95% (green) of its energy. Upper right, 20/30 Snellen E. Bottom left, aberrated Snellen E. Bottom right, left panel, optotype blurred by a circular kernel with the same diameter as the 50%-capturing circle at fixation. Bottom right, peripheral image of the apartment wall before and after blurring by the highly-aberrated off-axis PSF.

### 2.3 Eye Model

Using the Berkeley wide-field eye models^1^ and custom Zemax software, we estimated blur magnitudes across an 80° *×* 50° swath of the retina for each 3D view of the scene. The eye model simulation assumed that the eye was a “typical emmetrope” (as defined in Hastings et al.^1^, Fig. 1C), that the light source was 555 nm, and that the entrance pupil size, as imaged through the optics, was 4 mm in diameter. Blur magnitude was measured as the diameter of the circle that captured 50% of the energy of the monochromatic PSF (centered at the chief ray, Fig. 1E,H), and was computed for many combinations of object depth, fixation depth, and pupil size, yielding an interpolant function that could be used to linearly reconstruct the blur field corresponding to an arbitrary 3D view of a scene (retinal light field, Fig. 1F). The range of simulated object and fixation depths was 0 to 6 D, and the set of pupil diameters was 2, 4, and 6 mm. Accommodation was implemented by changes in lens curvature (and other eye model parameters) that best matched wavefront measurements collected during human accommodation (see Hastings et al.^1^ for details), thereby implicitly capturing accommodative leads and lags (Fig. 1G, left panel). Therefore, the model assumes a mild degree of presbyopia, and so when fixation depth was close, the accommodative state of the eye lagged slightly. For each view encountered by the virtual observer, we estimated the blur circle diameter for each field position using the interpolant function, and in the end, obtained a distribution of blur circle diameters for every position in the 80 *×* 50° retinal field across viewpoints (Fig. 1E). The simulation did not capture accommodation-linked pupil constrictions. These are quite small, but worth investigating in future extensions. We assumed the world was monochromatic (555 nm), even though real scenes are polychromatic and LCA strongly affects blur magnitudes. We found this to be justified for our particular goal of estimating the average blur field because comparisons showed that the encircled energy of a polychromatic PSF and the 555 nm PSF were very similar, and that 555 nm is near the most in-focus wavelength (i.e., centered in the LCA curve).

### 2.4 Image Simulations

To visualize how images are formed through the optics of the eye model or through the statistically average blur field (Fig. 1F,H), we took two different approaches. To visualize how an optotype is imaged through the central or peripheral optics, we computed the monochromatic point spread function (PSF) for 555 nm light at a specific field position and convolved it with a 20/30 Snellen E, for each RGB channel separately, performing reflective boundary handling. This procedure was justified because we were interested in the coarse level of blur subjectively for visualization purposes (Fig. 1H). To visualize the retinal image of the apartment scene during far and near work and across different pupil sizes, we used the image simulation tool in Zemax, which correctly incorporates LCA. For the composite images of near work (with the hand and phone), we superimposed two image simulations: one with the phone and hand, and one with the background image of the apartment, both with focus at 3.3 D, but the object depth at 3.3 D in the former and 0.67 D (the average object depth in our far work simulation) in the latter (Fig. 2). Blur magnitude is essentially asymptotic below 0.67 D due to dioptric space, so this planar assumption seemed justified because this composite image was for visualization purposes. Similarly, for the far work image simulation (Fig. 1F), blur circle diameters were nearly indistinguishable for the relaxed eye and median far work fixation/object depth (Fig. 2A-B), so we visualized the whole scene through an unaccommodated eye.

**Figure 2:**
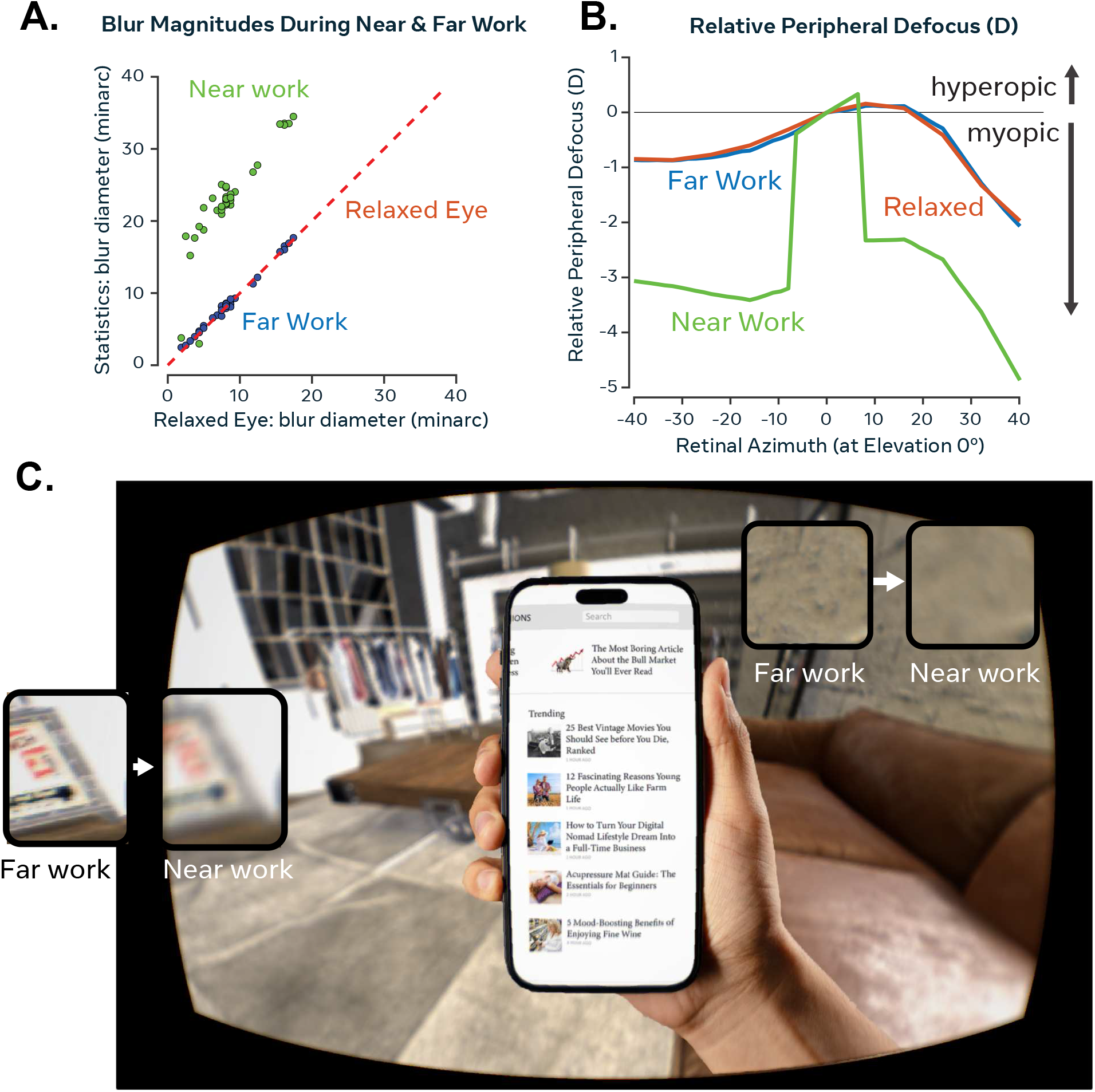
Near work induces extensive relative peripheral myopic defocus. **A.** Blur magnitudes are matched for far work and a relaxed eye, but 2–3*×* larger during near work. Average blur circle diameter (50% encircled energy) across the depth statistics encountered in far work and near work simulations versus for a “relaxed eye reference,” i.e., an unaccommodated eye looking at a frontoparallel plane at optical infinity. Each circle, one of 35 field locations depicted in Fig. 1E. Red dashed line, unity line depicting relaxed eye reference. **B.** Relative peripheral defocus across a horizontal slice of the retina at 0° elevation, estimated using paraxial curvature matching method (based on defocus and higher-order spherical Zernike coefficients).^49,50^ For a relaxed eye, the refractive state of the typical emmetrope eye model exhibits nasal-temporal asymmetries and moderate relative peripheral myopia. Far work does not alter this baseline profile. Near work induces 2–5 D of relative peripheral myopic defocus, when the fixation distance is 3.33 D and background objects now image behind the retina. **C.** Image simulation of aberrated retinal image of apartment scene during near work. Insets depict aberrated peripheral image during far work (reproduced from Fig. 1F) versus near work.

### 2.5 Simulating Near Work and Varying Light Levels

To simulate near work, we assumed that the observer was looking at a smartphone of realistic size and distance, while experiencing identical views of the environment in the background as specified in the active observer model. The phone was 14.7 *×* 7.2 cm or 26° *×* 13.4° in size, centered at fixation, at a distance of 3.33 D (30 cm) (Fig. 2C). So the focal distance of the eye (lagged accommodation response) was *∼*3 D.

To simulate the effects of indoor vs. outdoor lighting, we repeated the simulation with a 2, 4, or 6 mm pupil (Fig. 3). This allowed us to map the limits of pupil sizes experienced outdoors (*∼*2 mm) and indoors (*∼*6 mm), as well as those in-between. We did not make any other assumptions about how illumination may affect image formation.

**Figure 3:**
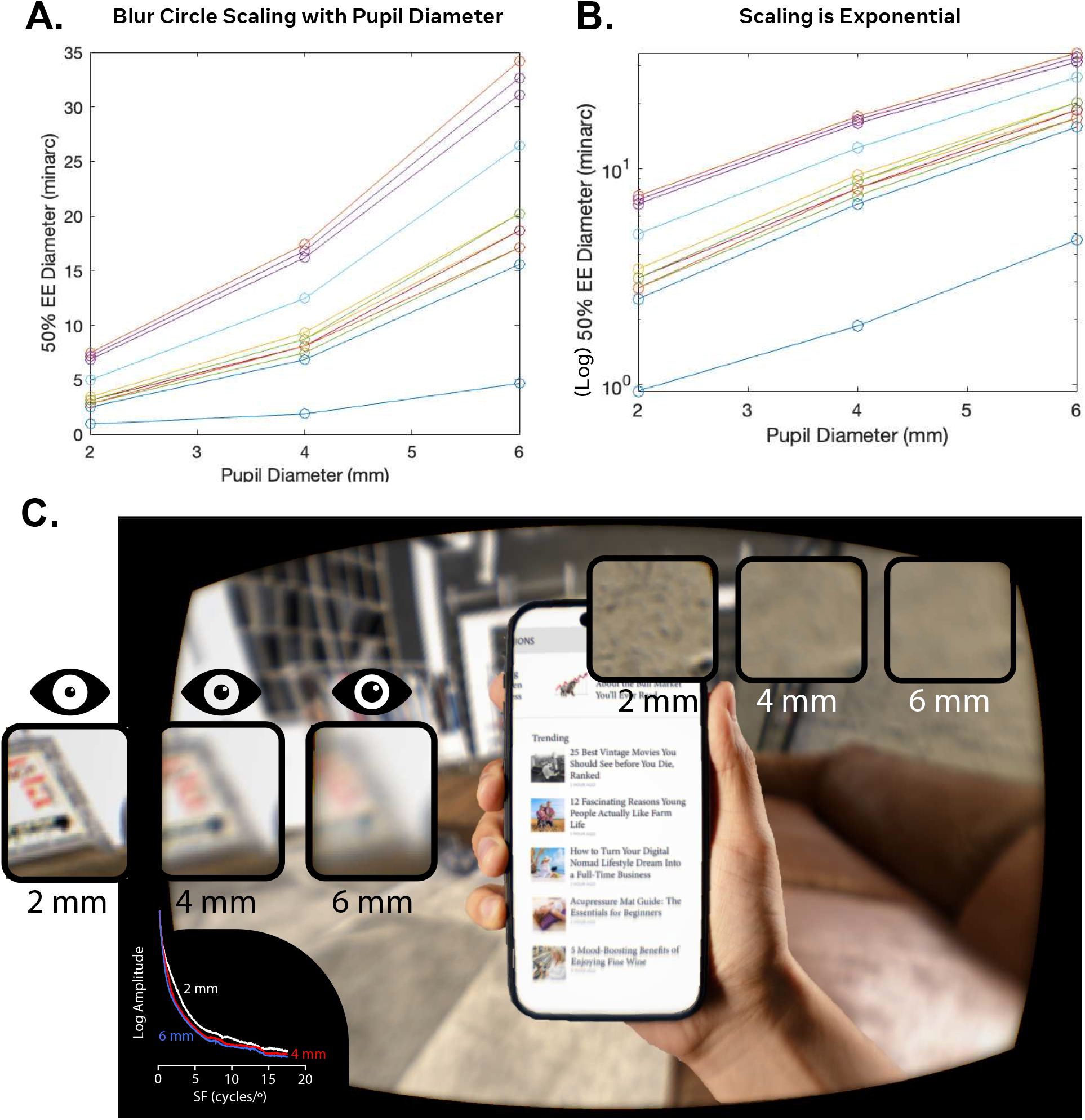
Low light levels exponentially increase blur diameter via pupil dilation. **A.** Blur circle diameter (50% encircled energy) in minarc for 2, 4, and 6 mm pupils, for 555 nm and an unaccommodated eye looking at a plane at optical infinity. Each colored curve corresponds to a specific field position. **B.** Data re-plotted with ordinate in log space. Blur scaling is now roughly log-linear for each field position, which means that the effect of pupil size on blur circle diameter is approximately exponential, where the exponent is fixed across field positions, but the intercept (i.e., baseline blurriness at that field position) varies. **C.** Near work at low light levels produces a highly blurred peripheral image. Image simulation depicts apartment scene through a 6 mm pupil, i.e., corresponding to very low light levels. Insets depict peripheral image through a 2, 4, and 6 mm pupil for ease of comparison, and the log amplitude spectrum for each, which demonstrates progressive contrast reduction across frequencies.

### 2.6 Bangerter Foil Simulation

It is challenging to directly simulate the effects of a specific diffuser filter on the retinal image in our framework without having a model of its physical properties. Instead, we simulated the subjective effects of a 0.4 strength Bangerter by defocusing the retinal image during far work (4 mm pupil) by an amount (1.73 D) that has been shown to produce the same subjective reduction in acuity as that specific diffuser in a psychophysical study^47^ (Fig. 4). We performed this by convolution with a circular kernel of diameter producing 1.73 D of blur (4 mm aperture), with reflective boundary handling. We felt this was a sensible approach because (1) objective measures of image degradation by Bangerter foils based on the image MTF under-blur relative to subjective measures like acuity, and (2) the aberrated image was very similar between the Bangerter aberrated image and the near work image, and contrast reductions were plausible. We measured image degradation (contrast reduction) as the percent reduction in the area under the radially-averaged (log-log) MTF of the resulting image between far work (4 mm) and the near work simulations at different pupil sizes and Bangerter foil (0.4).

**Figure 4:**
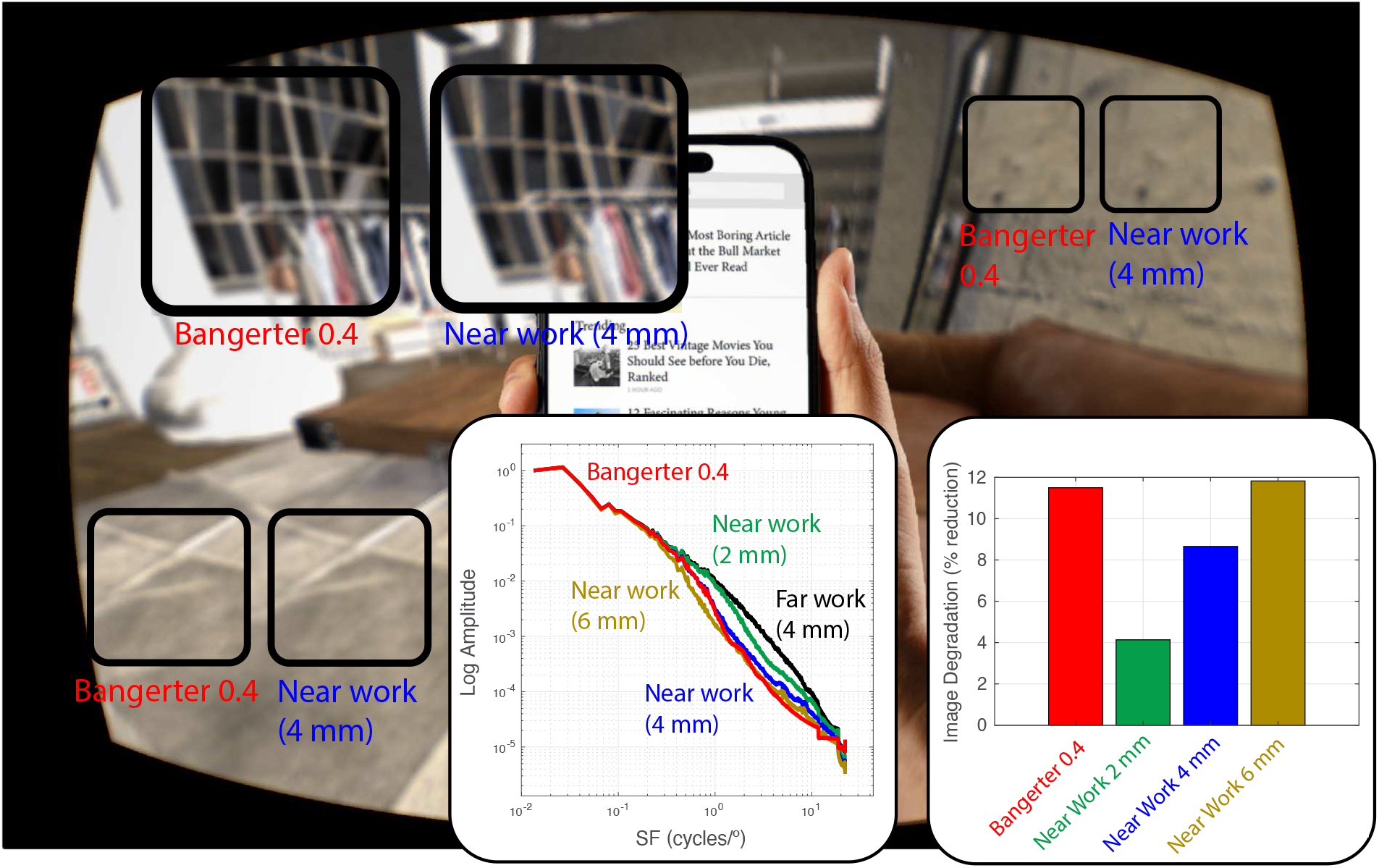
Peripheral contrast reductions induced by near work at low light levels and a Bangerter foil (0.4 strength) are roughly matched. Image simulation depicts example apartment view during near work through 4 mm pupil (reproduced from Fig. 2), against insets showing scene (during far work, 4 mm pupil) through a simulated 0.4 Bangerter foil (see Methods). Left inset, radially-averaged log amplitude spectrum against log frequency for example view. Black curve, far work (4 mm). Blue, near work (4 mm). Green, near work (2 mm). Red, Bangerter 0.4. Right, image degradation measured as percent reduction in area under the log amplitude spectrum with respect to far work (4 mm) reference spectrum, following Perez et al.^47^

### 2.7 Virtual Reality and Eye Tracking Experiment

To rule out the possibility that our findings are specific to the head and eye movements selected in our active observer model (which are not reward- and goal-driven by objects in the environment), we performed a validation experiment to collect real measurements of human fixation depths and head-centered depth maps during natural behavior. To avoid the small field-of-view imposed by camera and eye box limits of mobile eye trackers, we used a virtual driving experience in the Quest 3 head-mounted display, augmented with eye tracking (Fig. 6). Three observers (3 males, 30–40 years old) completed 10 minutes of the driving experience, which was implemented in Unity by modifying a comprehensive driving simulation Unity asset.^48^ The game required driving within a speed band (30–110 mph), so observers had to periodically check the speedometer on the dashboard (and therefore make near fixations as in real driving). A custom-integrated binocular eye tracker, Tobii Crystal (Tobii AB, Sweden; sampling rate, 240 Hz), was used to estimate 3D gaze position (vergence depth). We followed the manufacturer-recommended custom calibration and validation procedure, and only accepted calibrations with an average angular root mean squared (RMS) error across targets of less than 4°, and vergence depth error of 0.4 D. Head-centered depth maps were saved out on each frame, along with angular estimates of eye position and vergence depth. To estimate the retinal blur field on each frame, we computed the retinocentric depth map by shifting the head-centered depth map to center at the estimated angular position of the fixation point and then ran that depth map through the eye model, using the estimated vergence depth as a direct estimate of the fixation depth of the eye. Pupil size was assumed to be 4 mm, to match our other simulations, although the average measured pupil size across the three observers was slightly larger — around 4.5 mm. All observers had normal or corrected-to-normal near and far visual acuity of 20/25, color vision, and Randot stereo acuity of at least 100 arc seconds. Monocular near and far acuity were 20/30 or above, ruling out substantial clinical astigmatism (which could affect blur sensitivity). Participants were also screened for presbyopia (beyond 40 cm) and convergence insufficiencies with near point of accommodation and near point of convergence tests. All study protocols were IRB approved. Informed consent was obtained from all participants after explanation of the nature and possible consequences of the study. The research was conducted in compliance with the Declaration of Helsinki.

It is important to note that the focal plane (virtual image distance) in current commercial head-mounted displays is fixed. Therefore, users attempt to fix accommodation while moving vergence independently (resulting in vergence-accommodation conflict). This is not an issue for our purposes because we used the vergence and depth map information to simulate the retinal image during real driving (when vergence and accommodation should go together in non-presbyopes), not to model the retinal image during this specific virtual reality experiment.

## 3 Results

To estimate the natural statistics of retinal image blur in everyday life, we situate an active, accommodating, wide-field emmetropic eye model within a realistic 3D virtual apartment. In a first experiment, we constrain the model’s fixation selection process with published fixation depth (Section 2.2) and pupil size statistics measured during everyday activities (Section 2.5). This allows us to simulate the effects of far work and near work, as well as low lighting vs. outdoor lighting on optical aberrations to understand possible links between the visual diet and eye growth regulation. We compare the retinal conjugate surface and depth-of-field of the eye model with the natural statistics of object and focal depths to understand the human eye’s imaging properties. In a follow-up human experiment, we validate our findings with real measurements of fixation depths and large-field egocentric depth maps captured during a driving simulation in a head-mounted display augmented with binocular eye tracking.

### 3.1 Using the Active Eye Model to Estimate the Statistics of Retinal Blur During Far Work

To estimate blur magnitudes across the retina, we run many views of the apartment scene obtained from different realistic vantage points (Fig. 1A-B) and constrained by measured statistics of fixation depth in everyday life (Fig. 1C) through the typical emmetrope eye model.^1^ For each scene view, the object depth at a given retinal location (Fig. 1D), fixation depth, and pupil size jointly determine the blur magnitude there (Fig. 1E, grey circles). Across all views, the average blur circle scales with eccentricity roughly isotropically (Fig. 1E, grey circles), such that the retinal image is blurrier in the periphery and sharper (although not completely clear) at fixation (Fig. 1F). The average blur circle diameter is 2.5 minarc at fixation (0,0)° and 17.5 minarc in the corner of the modeled field (40,25)°.

At fixation (unity line in Fig. 1G), blur is primarily limited by defocus and higher order spherical aberrations, such that an object in front of or behind the focal distance of the eye will become blurrier roughly symmetrically (Fig. 1G, left panel). At fixation, objects are imaged nearly equally in focus regardless of their depth, due to accommodation. In the periphery, blur is dominated by off-axis aberrations including coma and astigmatism, creating very wide depth-of-field that creates nearly equally out-of-focus images of objects regardless of their depth and the eye’s fixation depth (depth-invariant imaging; Fig. 1G, right panel). At fixation (0,0)°, the PSF of the typical emmetrope eye has a punctate shape, and is primarily limited by defocus (Fig. 1H, left panel). The blur circle (50% encircled energy) is 2.5 minarc in diameter, and yields a reasonably focused image of a 20/30 Snellen E. Blurring the optotype by a circular kernel of the same diameter yields an image that is slightly more diffuse in appearance, but comparable in subjective resolution. At the corner of the field (40,25)°, the PSF is much larger and has a stereotypical coma-dominated shape (Fig. 1H, right panel). The blur circle is 17.5 minarc in diameter, and yields a phase-shifted, diffuse, and ultimately unrecognizable 20/30 Snellen E optotype. The roughly isotropic shape of the blur field is primarily due to the progressive predominance of these off-axis aberrations with increasing eccentricity.

### 3.2 Far Work Induces Retinal Blur That Is Mostly Indistinguishable from Retinal Blur of an Unaccommodated Eye

Due to very wide peripheral depth-of-field during far work, the average amount of blur across views is essentially identical to that of an unaccommodated (relaxed) eye (Fig. 2A). This tolerance holds not only for the average blur field, but also across the vast majority of individual views of the scene and their associated depth statistics (Fig. 1G, left panel). The profile of relative peripheral defocus is correspondingly similar during far work on average and for the unaccommodated eye (Fig. 2A).

### 3.3 Near Work Induces Relative Peripheral Myopic Defocus

We simulate near work in the apartment environment by having the virtual observer hold a smart-phone of realistic size (iPhone 14 dimensions: 14.7 *×* 7.2 cm; 26° *×* 13.4°) and distance (3.33 D), while experiencing otherwise identical backgrounds. Across views, the average peripheral blur circle diameter increased by 2–3*×* (versus for far work; Fig. 2A). This is due to a large degree of relative peripheral myopic defocus (2–5 D) across the peripheral retina, i.e., when attempting to focus at the phone, the background now images in front of the retina (Fig. 2B).

### 3.4 Blur Scales Exponentially with Pupil Size

In the well-known geometric approximation of defocus (which is only applicable to central vision), the diameter of the blur circle in radians scales linearly with pupil diameter. In the periphery, off-axis aberrations including astigmatism and coma are well-known to scale supralinearly with pupil diameter. When all aberrations in the eye model are considered in tandem for an eye focusing at optical infinity, the blur circle scales approximately exponentially in diameter with pupil diameter in the periphery (Fig. 3A-B). Outdoors during daylight, pupil diameter can range from 1.4 to 3 mm (average, 2.4 mm),^51^ and higher (*∼*4 mm) during cloudy weather.^52^ Indoors, light levels are lower due to the limited power of artificial light sources, and pupil size accordingly ranges from 3–6 mm depending on lighting and observer age (pupil size shrinks in old age).^51,52^ In our simulation, the blur circle is therefore approximately 2–3*×* larger in diameter for indoor vs. outdoor lighting. This increased blurring is clearly noticeable in an image simulation of near work at different pupil sizes (light levels), giving rise to reductions in image contrast at mid and high spatial frequencies (Fig. 3C).

### 3.5 Near Work at Low Light Levels Produces Peripheral Contrast Reductions Comparable to Moderate Diffusing Filters

The lack of mid and high spatial frequency detail in man-made, modern environments has been proposed as a possible environmental source of form deprivation in development.^33^ Here, we investigate whether form deprivation may also be a natural consequence of retinal blur by quantifying the contrast reductions observed during near work in indoor scenes and light levels in the peripheral retina due to the eye’s optics. We simulate the effects of a moderate diffuser filter (Bangerter foil, 0.4 strength) on the retinal image (see Methods) and quantify the total contrast reduction during near work and for the Bangerter foil relative to far work in the same scene. Contrast reduction is quantified as the percent reduction in the area under the MTF relative to far work through a 4 mm pupil. Near work through a 4 or 6 mm pupil both induce similar amounts of blur to a 0.4 Bangerter foil, suggesting that they may induce a degree of image degradation comparable to levels known to affect eye growth in animal models. The image simulation in Fig. 4 shows how subjectively similar the retinal image is during 4 mm near work and through the Bangerter foil.

### 3.6 Peripheral Blur Is Distance-Invariant with Far Work

For the model observer performing far work in the apartment scene, the average fixation depth was 0.67 D or 1.5 m (Fig. 1C, right subpanel), with a good deal of variability across fixations (Fig. 1G, right subpanel). Likewise, the typical depth range for a single view of the apartment is large (e.g., *>* 2.8 D or *∼*5 m; Fig. 1D), such that many objects fall in front or behind the fixation point. And across many views, in the periphery, object depths vary widely between 0 and 6 D (Fig. 1G, right subpanel). However, the blur circle diameter at (40,25)°, for example, is virtually unchanged (plus/minus a few minarc) regardless of whether an object is far or near, when the eye is focused beyond *∼*1 m, i.e., during far work. That is, the peripheral image during far work is roughly equally out-of-focus, or depth-tolerant, regardless of object distance.

There are two interacting phenomena for this depth-tolerance. First, off-axis aberrations like coma and astigmatism predominate in the periphery (Fig. 1H, right subpanel), and these aberrations are mostly invariant to the depth of a light source of matched intensity. In other words, depth-of-field is large in the periphery (Fig. 5, top), and nearly all objects (95%) in our simulation are within a conservative measure of the depth-of-field (e.g., within +2 minarc in the blur circle).

**Figure 5:**
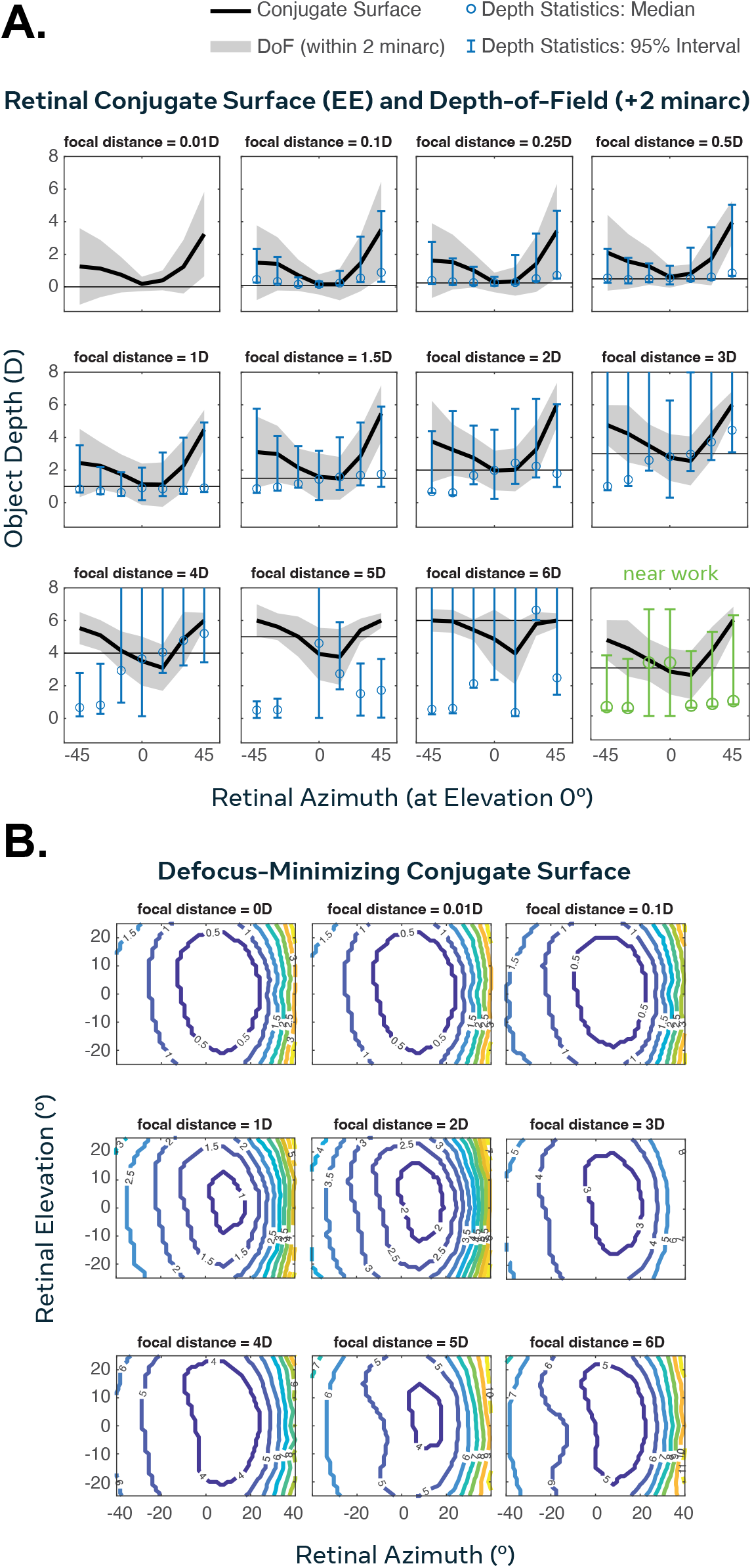
Retinal conjugate surfaces based on the 50% encircled energy blur circle or based on estimated defocus alone, both for the typical emmetrope eye model. **A.** The human eye achieves nearly depth-invariant imaging during far work. Black curve, retinal conjugate surface across the horizontal meridian of retina (elevation 0°) obtained by finding the depth minimizing the blur circle (50% encircled energy) for a given fixation distance (thin horizontal reference line). Grey surface, depth-of-field around conjugate surface (yielding +2 minarc of additional blur). Blue circle, median object depth for far work apartment simulation across horizontal median of retina. Error bar, central 95% of the distribution of object depths. Note, the apartment contains no object depths beyond 0.01 D, and very few below 5 D. Green curve, near work simulation, with fixation depth of 3.33 D. Note how most object depths are behind conjugate surface and depth-of-field in the peripheral retina. **B.** Contours of 3D defocus-minimizing conjugate surface across 40° by 25° field. Defocus estimated using paraxial method accounting for higher order spherical aberrations, using Zernike coefficients of typical emmetrope eye model fixating on a range of distances between 0 and 6 D.^1,49,50^ Contours depict conjugate surface in diopters. Note the extensive curvature, which matches the curvature of the conjugate surface in panel A.

Second, the retinal conjugate surface — the set of points in the world that is most in focus at each part of the retina — is curved towards the eye in the periphery (we estimate the conjugate surface by finding the object depth that produces the minimum 50% encircled energy blur circle for an eye attempting to focus at a number of different fixation distances). From a bird’s eye view, the conjugate surface is more curved toward the face than one might intuit, spanning almost 4 D from fixation to 40° of eccentricity. Because of this, the most in-focus object at (40,25)° is actually just a few centimeters away from the eye, and the origin of the asymmetry in Fig. 1G. This curvature (the ridge of the conjugate surface) is due to the geometry of defocus (Fig. 5, bottom), and the shape of the human globe. Therefore, this curvature is likely not an evolutionary adaptation to the comparatively mild degree of enclosure observed in environmental depth maps (Fig. 5, top).^8^ Note that the conjugate surface is not equally in focus across eccentricity. It also has large depth-of-field around it, and therefore is better conceived of as a very wide plateau, such that even the terminology “best-in-focus” is misleading. Near work breaks this depth-invariance by moving the background off this wide conjugate surface (Fig. 5, top, “near work”).

### 3.7 Human Fixation Depth and Retinocentric Depth Map Measurements During Virtual Driving Validate Peripheral Depth-Tolerance

We obtained measurements of human fixation depths (from convergence distance estimates) and retinocentric depth maps during simulated driving in a Quest 3 headset augmented with binocular eye tracking (Fig. 6A), and ran these through the typical emmetrope eye model (see Methods), in order to validate our simulation findings. We observed depth-tolerant imaging during driving across the retina, arising from the observation that virtual driving mostly constituted far work (the majority of measured fixation depths were greater than 5 m, Fig. 6B). The median fixation depth and object depth (at 0°,25°) were correspondingly below 1 D, as the observer looked at the road and other objects outside the window (Fig. 6C). Interestingly, in the far periphery, the blur circle was sometimes actually up to 5 minarc smaller in diameter during driving than for an unaccommodated reference eye looking at a plane at optical infinity. This counter-intuitive result would be explained by the extreme curvature of the conjugate surface, such that objects inside of the car should be slightly more in focus in the periphery than objects outside the car (Fig. 1G, Fig. 5A) — and that the peripheral image was often composed of the dashboard and other parts of the car’s interior. This experiment using human fixation and object depths (obtained during unrestricted eye and head movements) suggests that our procedure is relatively robust to the artificial head/eye movements and placement assumptions made in our far work active observer simulation.

**Figure 6:**
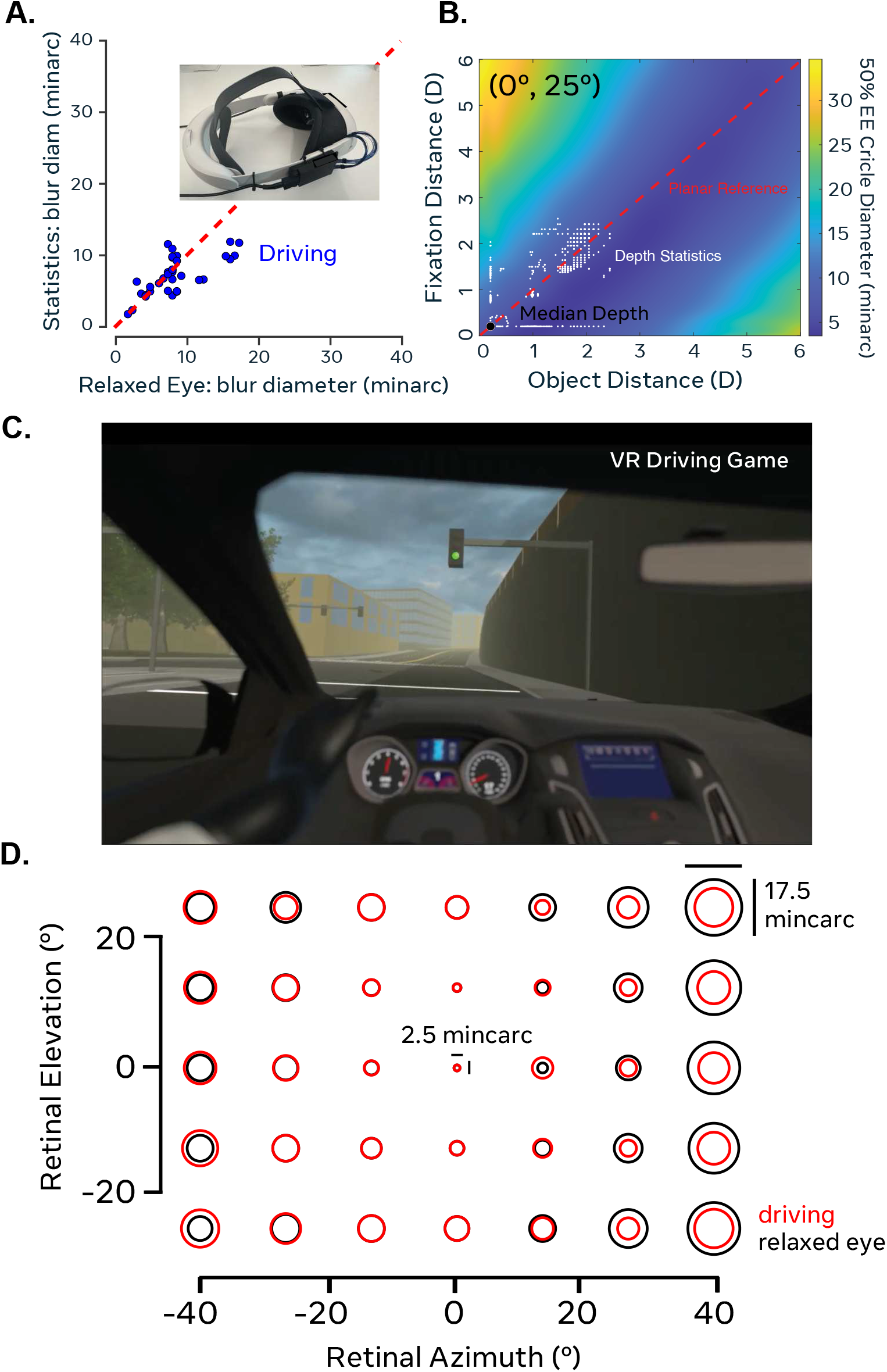
Measurements of eye and head movements during virtual driving experiment validate depth-tolerance of peripheral imaging during far work. **A.** Average blur circle diameter integrated across 3D scene and fixation depth statistics during virtual driving from a representative observer (other observers were very similar) vs. relaxed eye reference for each field position. Inset, Quest 3 head-mounted display augmented with Tobii binocular eye tracker. **B.** Depth-of-field at (0,25)° retinal location. Same format as Fig. 1G, with data from representative observer. Note that depth statistics lie in roughly level basin function, i.e., depth-tolerance. Clipped values on edges are at or beyond 0.25 D, so clipping errors have negligible impact on results. **C.** In-game view of driving simulation. **D.** Average blur circle during driving across field for representative observer, versus relaxed eye reference.

### 3.8 Popular Geometric Approximation of Defocus Mis-estimates Blur Magnitude

The common geometric approximation of defocus is useful for optometrists and vision scientists to gain intuition about image formation in central vision, where image quality is primarily limited by defocus. However, this approximation has been used to model blur outside of central vision in a number of papers, and has even been applied in the periphery.^42,53,54^ The approximation is

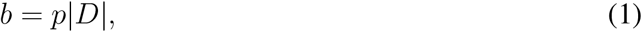

where *b* is blur circle diameter in radians arising from defocus, *p* is pupil diameter in meters, and *D* is the relative depth between the focal distance and an image-forming object. Applying such a model outside of fixation is incorrect for a number of reasons (off-axis aberrations and their induced depth-of-field, geometry of defocus and conjugate surface), but the accuracy of this approximation is not well-characterized. Sprague et al.^42^ used the geometric approximation of defocus to estimate the blur field across the central 20° of the retina across four everyday tasks (ordering coffee, making a sandwich, walking inside, and walking outside), which included a mix of both near and far work. We re-analyzed their data, expressing it in units of 50% EE diameter. As expected, image focus is unrealistically sharp in central vision for the approximation vs. more realistic optics (Fig. 7A). In the parafovea, blur is also underestimated by the geometric model. The reasons for this are two-fold. The conjugate surface of the geometric model is a frontoparallel plane, if one interprets depth in the classic sense in computer vision as depth along the line-of-sight (Fig. 7B). The depth-of-field of the geometric approximation is also much narrower than for the eye model, even at fixation (Fig. 7B, compare with Fig. 1G). A direct comparison of blur circle diameter (50% EE) between the geometric approximation and the eye model, given the same input depth statistics from the far work simulation reveal that the geometric model would grossly underestimate blur magnitudes across the retina (Fig. 7C). However, the geometric model of defocus does not always underestimate retinal blur magnitude in general. The mis-estimation is complex and the sign of the bias depends on the field position eccentricity and the focal distance of the eye (and thereby on relative depth as well). When relative distance is low, it can often overestimate blur magnitude (which was noted by Sprague et al. as well).^42^

**Figure 7:**
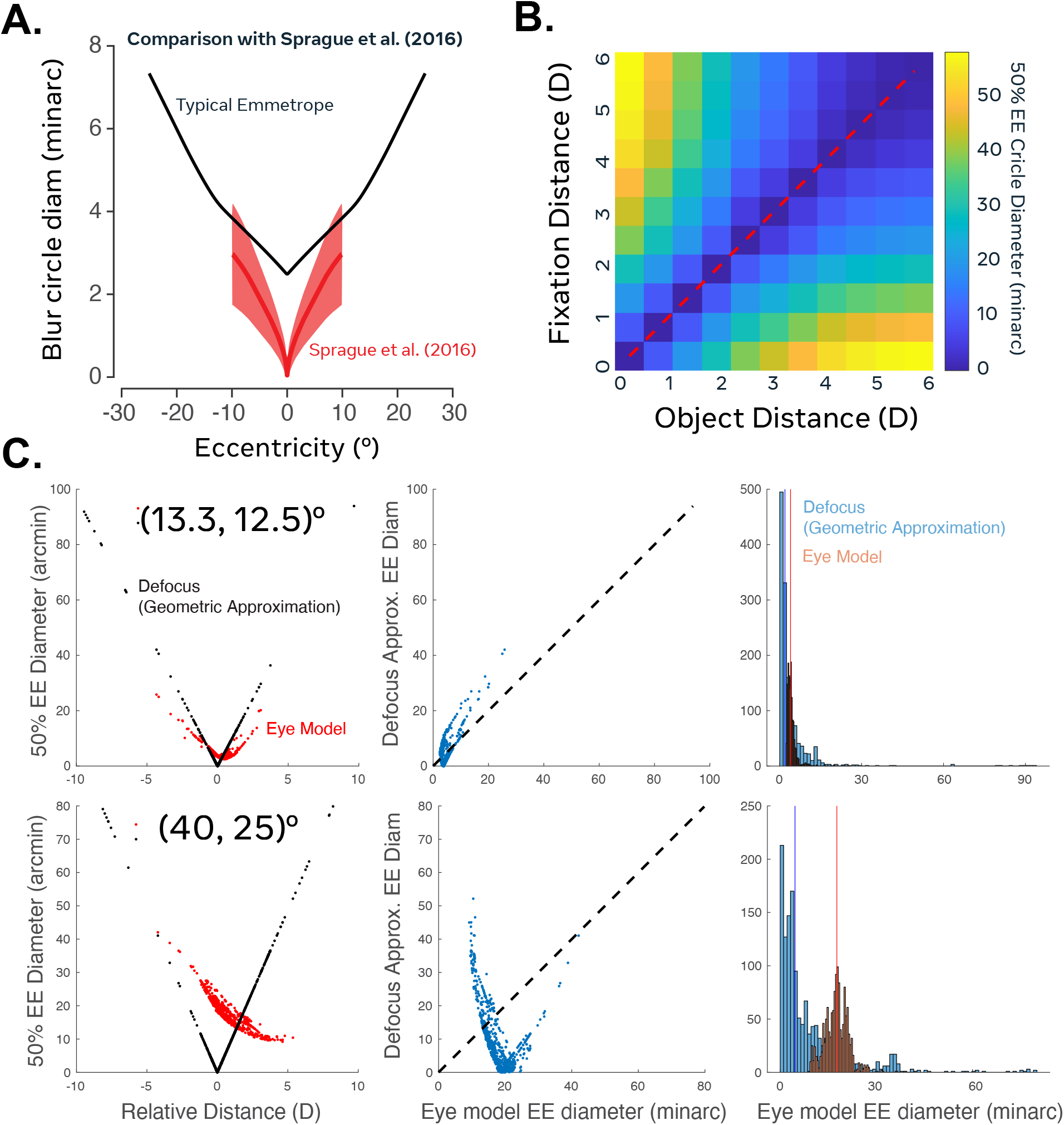
Comparison of blur magnitude across the central 20° for the geometric approximation of defocus employed in Sprague et al.^42^ vs. the typical emmetrope eye model,^1^ depth-of-field of defocus approximation, and direct comparison of blur magnitudes from eye model vs. geometric approximation given the far work depth statistics from our active observer model. **A.** Red curve, radial average of the average 50% blur circle diameter across the four tasks in Sprague et al., data re-analyzed. Error surface, SEM across the four tasks. Black line, 50% radial average of blur circle diameter for our far work simulation. **B.** Depth-of-field for geometric approximation of defocus is much narrower than for typical emmetrope eye model at fixation (compare with Fig. 1G). Also note the difference in colormap limit — the maximum blur circle from the approximation of defocus is larger than for the eye model in central vision (which is mostly limited by defocus). The imputed conjugate surface will be a frontoparallel plane if you apply this approximation (appropriate for central vision) in the periphery, as Sprague et al. and many past papers did. **C.** Direct comparison of blur magnitudes between eye model and geometric approximation of defocus given the same input depth statistics. Left panel, 50% EE blur circle diameter (minarc) as a function of relative distance between fixation and object depth for the typical emmetrope eye model (black) and geometric approximation of defocus (red) at mid-peripheral (13.3, 12.5)° and far-peripheral (40, 25)° retinal locations. Middle panel, scatterplot of blur circle diameter for the two against each other. Right panel, histogram of blur magnitudes across views of scene for each. Vertical lines, median blur circle diameter.

## 4 Discussion

We introduced active eye modeling, a computational platform combining wide-field accommodating eye models,^1^ realistic three-dimensional virtual environments,^44^ and naturalistic gaze patterns constrained by human data^42,45^, to estimate retinal image statistics under conditions representative of modern visual experience. Our simulations yield three principal findings. First, far work (*∼*1 m or greater) produces depth-tolerant peripheral imaging: off-axis aberrations create such wide peripheral depth-of-field that the retinal blur pattern is essentially indistinguishable from that of a relaxed, unaccommodated eye. Second, near work under indoor lighting substantially increases peripheral blur and reduces contrast at mid and high spatial frequencies. Third, the magnitude of this optical contrast reduction is comparable to that produced by a moderate diffusing filter (Bangerter 0.4 strength), a degree of image degradation that affects eye growth when imposed experimentally in animal models.^13,14^ A consistency check using eye-tracked virtual driving confirmed the depth-tolerance of peripheral imaging during far work.

### 4.1 Near Work Induces Peripheral Myopic Defocus

Our simulations reveal that near work in typical indoor environments induces extensive relative peripheral myopic (not hyperopic) defocus: when the eye accommodates to a near target, background objects at greater distances image in front of the retina. This is consistent with the geometry of near work and provides quantitative confirmation complementing recent time-of-flight measurements of the dioptric landscape during near-work activities.^8^ The magnitude of this peripheral myopic defocus (2–5 D) far exceeds the relatively small changes in defocus expected from accommodative lags in young adults,^11^ although the two operate at different retinal eccentricities and may engage different components of the growth-regulation system. This result illustrates how the peripheral defocus pattern during near work depends critically on the full dioptric landscape of the environment, not just the fixation distance.

The myopic direction of peripheral defocus during simulated near work does not readily support the hypothesis that near work drives myopia specifically via peripheral hyperopic defocus causing eye elongation. This is consistent with the growing recognition that defocus direction alone may not be the critical variable: lenses inducing either sign of peripheral defocus can slow myopia progression in clinical trials.^10,27^ Longitudinal studies show that relative peripheral refraction measured clinically does not reliably predict who will develop myopia^40,41^ and may reflect a consequence rather than a cause of refractive error.^26,39^ Standard clinical measurements provide only a static snapshot at optical infinity and a few peripheral locations. The actual retinal signals experienced during near work at indoor light levels, as our simulations illustrate, may differ substantially. Understanding how optical interventions interact with the defocus patterns generated by the wearer’s real visual diet, which this platform can simulate, may help resolve apparent contradictions in the efficacy of different lens designs.

### 4.2 Peripheral Blur and Contrast Reduction

Near work at typical indoor light levels (pupil diameter *∼*4–6 mm) produces peripheral optical contrast reductions at mid and high spatial frequencies that are comparable in magnitude to those of a 0.4-strength Bangerter diffuser,^47^ as quantified by the area under the radially-averaged MTF. It is important to note that the physical mechanisms of image degradation differ: near-work blur arises from aberration-limited defocus producing asymmetric, phase-shifting PSFs, whereas Bangerter foils produce broadband forward scattering with a more symmetric halo. Whether these different degradation patterns produce equivalent biological effects on emmetropization signaling is an open empirical question. Nonetheless, the predicted magnitude of contrast reduction is notable, as diffusing filters of this strength reliably affect eye growth in animal models.^13,14^

This connects to the proposal that man-made environments may lack the high spatial frequency content needed for normal emmetropization.^33^ Notably, man-made environments are also not scale invariant, so halving viewing distance indoors further reduces middle spatial frequency contrast by approximately 35%.^34^ Our results suggest that the eye’s own optics, interacting with near work and dim lighting, may produce a functionally similar degree of contrast reduction even in environments with abundant fine detail. If such blur constitutes a graded stimulus, an assumption that remains to be tested, then chronic exposure during near work at indoor light levels could contribute to a dose-dependent effect on eye growth.

However, this finding must be interpreted in light of evidence that mild peripheral contrast reduction via DOT lenses can actually slow myopia progression.^29,31^ Two observations (strong reduction promotes elongation; mild reduction may inhibit it) suggest the relationship between peripheral contrast and eye growth may be non-monotonic, but the mechanisms may also differ fundamentally: DOT lenses may act through spatial frequency-specific contrast changes, phase disruption, scattering-induced shifts in the peripheral image shell, or effects on accommodation, rather than through bulk contrast reduction alone.^24,28^ The involvement of retinal ON and OFF contrast pathways in refractive development^29,30^ adds further complexity.

### 4.3 Depth-Tolerant Peripheral Imaging During Far Work

During far work, the peripheral retina exhibits depth-tolerant imaging: blur magnitude changes minimally regardless of object distance within the range of naturally occurring depths, for two interacting reasons. Off-axis aberrations (primarily coma and astigmatism) dominate peripheral image quality and are largely insensitive to object distance, creating wide depth-of-field. Additionally, the retinal conjugate surface curves substantially toward the observer in the periphery due to the geometry of defocus and globe shape, creating a wide depth-of-field plateau within which virtually all naturally occurring object depths fall during distance viewing. Our consistency check during virtual driving confirmed this prediction with measured human fixation and depth statistics.

This depth-tolerance could represent a functionally relevant feature of far-work optics: the peripheral retina receives a consistent optical signal regardless of environmental depth variability, which may support stable emmetropization, though whether peripheral image stability per se contributes to emmetropization remains to be determined. Near work breaks this depth-tolerance by moving background objects off the wide conjugate surface, producing the large peripheral blur documented above. The interaction with pupil size amplifies the effect: far work outdoors (small pupil, distant fixation) produces the sharpest peripheral images, while near work indoors (large pupil, near fixation) produces the most degraded. Our finding that blur scales approximately exponentially with pupil diameter in the periphery (documented in the Results) provides an optical mechanism through which time indoors could contribute to increased peripheral blur, complementing dopamine-mediated mechanisms.^36–38^

### 4.4 Future Directions

Several extensions of this platform would be valuable. First, simulating retinal image statistics through myopic eye models^1^ could reveal how peripheral defocus patterns differ from those of emmetropic eyes, testing whether the optical conditions we identify contribute differently to myopia onset versus progression. Second, the current platform simulates individual static views; a natural extension is to simulate temporal sequences capturing dynamic variation in blur and contrast. This is relevant because temporal luminance modulation at different frequencies can differentially affect eye growth,^55–60^ and dynamic contrast variation — particularly differential effects on positive and negative contrast through ON and OFF pathways — remains largely unexplored in the context of emmetropization. Third, the platform could simulate how specific optical interventions (e.g., micro-lenses technology such as HAL or DIMS, bifocal lenses, and/or orthokeratology) interact with realistic environmental depth structure during everyday activities, going beyond bench-top optical characterization. Finally, because the framework accepts real-world inputs, i.e., depth maps from camera or time-of-flight sensors, measured gaze and vergence depth, and video that can be optically degraded, it can reconstruct the retinal image directly from data captured during everyday behavior, not only from simulated environments. Integration with emerging wearable sensing technologies that measure gaze, light exposure, and viewing distance would enable model validation and increasingly realistic, individualized estimates of the visual diet, creating a virtuous cycle between measurement and prediction.

Beyond refractive development, the optics form the earliest information bottleneck in visual processing, and our estimates of the natural blur field across eccentricity may help constrain theories of retinal and cortical organization during development.^61^ This illustrates the platform’s broader utility for questions about peripheral vision and visual development.

### 4.5 Limitations

Our simulations assume monochromatic light (555 nm), although longitudinal chromatic aberration strongly affects blur.^1^ We verified that encircled energy of the polychromatic and 555 nm PSFs are similar for the purpose of estimating the average blur field, but this precludes conclusions about chromatic mechanisms of emmetropization.^22,62^ We did not model accommodation-linked pupil constriction; this effect (typically 1–2 mm during strong convergence) would partially counteract peripheral blur during near work, though given the approximately exponential blur-pupil relationship, the optical consequences of this constriction are modest relative to the large contrast reductions we document. The “typical emmetrope” eye model^1^ represents one pattern of nasal-temporal asymmetry and mild presbyopia. Because the developing eye differs from the adult emmetropic eye in axial length, corneal power, crystalline lens structure, and aberration profile, the quantitative blur and defocus patterns reported here should be interpreted as illustrative of the optical principles rather than as direct estimates of the pediatric retinal image;^63^ extending the platform to age-appropriate eye models is a natural next step. The shape of the retinal conjugate surface we observe is determined by the geometry of defocus and the globe shape, rather than being an adaptation to environmental depth statistics; this is consistent with evidence that the vertical horopter is not adaptable to experience.^64^ Our virtual apartment, while realistic, represents one indoor environment; the platform generalizes to other environments, and the depth-tolerance we observe during far work suggests the blur estimates are relatively robust to environmental variation. The eye-tracked VR consistency check is limited by N=3 (all male, 30–40 years old), a single far-work task (driving), vergence estimation noise, and vergence-accommodation conflict in the head-mounted display. Importantly, the near-work/low-light predictions, one of our central findings, were not empirically validated and remain to be tested.

In summary, active eye modeling provides a flexible computational platform for investigating how the visual diet shapes retinal image formation. Applied to common modern visual behaviors, the platform identifies optical conditions during indoor near work — substantial peripheral blur and contrast reduction arising from the interaction of near accommodation, dim lighting, and the eye’s own off-axis aberrations — that warrant empirical investigation as factors in refractive development. More broadly, because the retinal image is the common input to all downstream vision, active eye modeling is a general tool for any problem that depends on egocentric visual statistics, including the design of displays, spectacles, and contact lenses, naturalistic psychophysics, and the study of peripheral vision, accommodation, and visual development, questions that are difficult to address with current measurement technology alone.

## Acknowledgments

Claude (Sonnet 4.5, Anthropic), Oct–Nov 2026, was used for LaTeX formatting, reference formatting, and copyediting of author-written text.

## Notes

### Competing Interest Statement

The authors are affiliated with Meta Platforms, though this study did not include any products or services related to the company. As this is not evaluating or promoting any products or services, we do not have any competing or financial interests to report.

